# Liquid-liquid phase separation of the chromosomal passenger complex drives parallel bundling of midzone microtubules

**DOI:** 10.1101/2022.01.07.475460

**Authors:** Ewa Niedzialkowska, Tan M. Truong, Luke A. Eldredge, Stefanie Redemann, Denis Chretien, P. Todd Stukenberg

**Affiliations:** Department of Biochemistry and Molecular Biology, University of Virginia, School of Medicine, Charlottesville, VA, USA; Department of Cell Biology, University of Virginia, School of Medicine, Charlottesville, VA, USA; Department of Molecular Physiology and Biological Physics, University of Virginia, School of Medicine, Charlottesville, VA, USA; Center for Membrane and Cell Physiology, University of Virginia, School of Medicine, Charlottesville, VA, USA; Univ Rennes, CNRS, IGDR (Institut de Génétique et Développement de Rennes) - UMR 6290, F-35000 Rennes, France

## Abstract

The spindle midzone is a dynamic structure that forms during anaphase, mediates chromosome segregation, and provides a signaling platform to position the cleavage furrow. The spindle midzone comprises two antiparallel bundles of microtubules (MTs) but the process of their formation is poorly understood. Here, we show that the Chromosomal Passenger Complex (CPC) undergoes liquid-liquid phase separation (LLPS) to generate parallel MT bundles *in vitro* when incubated with free tubulin and GTP. MT bundles emerge from CPC droplets with protruding minus-ends that then grow into long, tapered MT structures. During this growth, the CPC in condensates apparently reorganize to coat and bundle the resulting MT structures. CPC mutants attenuated for LLPS or MT binding prevented the generation of parallel MT bundles *in vitro* and reduced the number of MTs present at spindle midzones in HeLa cells. Our data uncovers a kinase-independent function of the CPC and provides models for how cells generate parallel-bundled MT structures that are important for the assembly of the mitotic spindle.

## Introduction

During anaphase, microtubule (MT) structures form in the central region of the spindle known as the spindle midzone (hereafter midzone), which comprises parallel-bundled MTs linked by antiparallel cross-linked plus ends to define the cytokinetic furrow at the cell cortex [1, 2]. Midzones have key functions in anaphase, such as roles in anaphase B movements of chromosomes by generating a force that pushes poles apart [3] and in generating a signal that defines the location of the cytokinetic furrow [4]. Antiparallel MTs are cross-linked at midzones by Protein Regulator of Cytokinesis 1 (PRC1) and capped at MT plus-ends by the kinesin Kif4a [5, 6], which both work with Cytoplasmic Linker-associated Proteins (CLASPs) in order to regulate the stabilization, dynamics, and formation of midzone MTs [7]. They localize to the central region of the midzone [8] along with a set of proteins required to generate the signal for the generation of the cytokinetic furrow known as the centralspindlin complex.

The Chromosomal Passenger Complex (CPC) is another regulator of late mitosis. The CPC shows dynamic localization throughout mitosis as it moves from inner centromeres to the midzone during the metaphase to anaphase transition [9, 10] to form and stabilize the spindle midzone in anaphase, and to regulate cytokinesis [11]. The CPC is a heterotetrameric assemblage of the scaffolding protein, INCENP, that binds and regulates the kinase Aurora B at its C-terminus, and binds to both the chromatin-binding Survivin and phase-separating Borealin via a coiled-coil at its N-terminus [12]. At later stages of cell division, the CPC is required for the generation of signals that define the cytokinetic furrow [13] and identify lagging anaphase chromosomes [11, 14, 15]. The CPC also contains at least two MT-binding sites: one within the N-terminus of Borealin [16] and the other a single α-helix (SAH) domain of INCENP [17]. It is yet unclear, however, whether the CPC functions are limited to regulating and localizing Aurora B kinase signaling or there exists additional roles pertaining to organizing microtubules in the midzone.

It has recently been shown that the centromere-targeting region of the CPC, which is composed of Survivin, Borealin, and the N-terminus of INCENP (hereafter referred to as the CEN domain), binds and bundles MTs *in vitro*. This property is thought to be mediated by the Borealin subunit, which contains an N-terminal MT-binding site [16] and drives liquid-liquid phase separation (LLPS) of the CEN domain [18]. The importance of the CPC LLPS and MT bundling activity is unclear, but the amount of the CPC in midzones is reduced in mutants deficient in CPC LLPS [18] and the complex did not bind to the central spindle or the midbody if mutated in the MT-binding region of Borealin [19].

Here, we investigate how properties of the CPC to both bind MTs and undergo LLPS contributes to formation of the spindle midzone. We first expressed recombinant CPC and demonstrate its ability to undergo LLPS *in vitro*. We then find that CPC condensates concentrate free tubulin and, in the presence of GTP, facilitate the formation of parallel MT bundles. These bundles emerge from CPC condensates with their minus ends protruding distally, which is the opposite polarity to other microtubule-organizing centers (MTOCs), such as centrosomes and Golgi apparatus. Midzone formation is deficient in cells expressing CPC mutants deficient in either MT-bundling or LLPS. Altogether, our data indicate that the CPC exists in a liquid-demixed state at the midzone and MT organizing activities emerge from the combination of its MT-binding and liquid-demixing properties.

## Results

### The chromosomal passenger complex undergoes LLPS *in vitro*

The *Xenopus laevis* CPC (Figure 1A,B) was expressed in *E.coli* and purified on nickel-nitrilotriacetic acid (Ni-NTA) resin via a 6xHIS tag fused to the N-terminus of INCENP. The purified complex migrated on size exclusion chromatography as a single peak under high salt conditions (Supplementary Figure 1A). Although the complex appears to have greater amounts of Borealin by Coomassie staining, that protein is highly enriched in K/R amino acids that bind Coomassie [20] and an equal stoichiometry of all four subunits was confirmed by mass spectroscopy analysis (Supplementary Figure 1B). A disordered region of Borealin (amino acid residues 139-160) is involved in LLPS of a subcomplex of the CPC [18]. Recombinant CPC undergoes LLPS *in vitro* in the presence of a crowding agent (7% PEG3350) or in a low salt buffer, as in these conditions, we observed the formation of spherical droplets characteristic of LLPS (Figure 1C). These droplets fused at early time points after inducing phase separation, demonstrating their liquid properties (Figure 1D). The CPC droplets formed at 1 μM, which is below the concentration that has been estimated for Aurora B at centromeres [21]. *In vitro* CPC droplets retained specific binding to histone H3 peptide phosphorylated on T3 and Aurora B kinase activity (Figure 1E, F).

**Figure 1.**
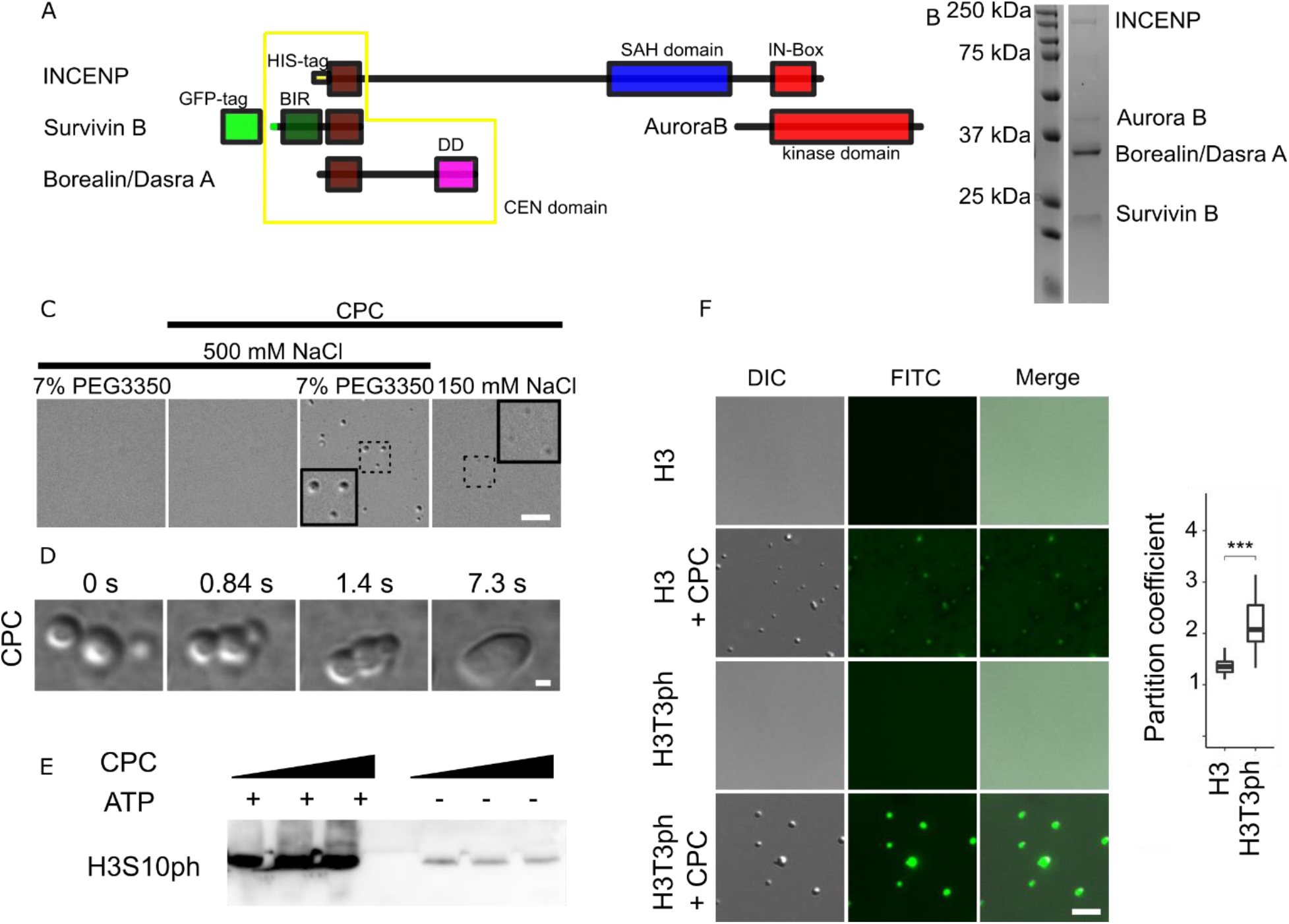
Recombinant CPC undergoes liquid-liquid phase separation. A. Scheme of the CPC; CEN: centromere-targeting domain; SAH: single α-helix domain; IN-Box: INCENP conserved box; DD - dimerization domain. B. SDS-PAGE gel with Coomassie staining of recombinant CPC C. Differential Interference Contrast (DIC) of CPC condensates; scale bar = 5 μm. Insets showed magnified views of dotted regions. D. DIC images showing fusion of the CPC coacervates; scale bar = 1 μm. E. Western blot confirms the phosphorylation of histone H3 Ser10 in the presence of 0.25 μM, 0.5 μM and 1 μM LLPS CPC and ATP. F. DIC and fluorescence images showing partitioning of histone H3T3ph (1-21)-FITC and H3T3 (1-21)-FITC into CPC condensates. Plot to the right shows calculated partition coefficient of H3T3ph (1-21)-FITC peptide (n=42) and H3T3 (1-21)-FITC peptide (n=42) into coacervates. Experiment was repeated twice; p-value = 2.2·10^-16^; scale bar = 5 μm. Statistical analysis was performed by applying Kolmogorov-Smirnov test, * = p<0.05, ** = p < 0.01, *** = p<0.001.

### The CPC nucleates and bundles MTs

In cells, MT nucleation is mostly driven by γ-tubulin ring complexes (γ-TuRC), which generates MTs of 13 protofilaments [22–24]. However, chromatin-driven spindle assembly pathways independent of γ-TuRC have been reported in meiotic systems [25], in plant cells that do not contain centrosomes [26], and on DNA-coated beads [27]. The CPC regulates assembly of chromatin-associated MTs by controlling the activity of the microtubule depolymerase, MCAK [28, 29] *in vivo*. However, it is unclear if the CPC alone is sufficient to directly regulate microtubules formation. We previously showed that α/β-tubulin dimers are enriched in condensates of a CPC subcomplex. Mixing these condensates with tubulin in the presence of GTP generated elongated MT structures, and paclitaxel stabilized MTs were bundled by this CPC subcomplex [18]. Here, we tested whether the full-length CPC could also concentrate free tubulin and generate microtubule structures. α/β-tubulin dimers were approximately five-fold enriched in CPC condensates (Figure 2A). As a control, GFP showed no preferable enrichment in condensates compared to the dilute phase (Supplementary Figure 2A). When CPC coacervates were incubated with free tubulin in the presence of GTP, elongated structures were observed within 10 minutes of incubation (Figure 2C). No MT structures were formed in the absence of GTP or CPC (Figure 2B; Supplementary Figure 2B) and GMpCpp polymerized MTs incubated with PEG3350 did not bundle (Supplementary Figure 2C). We imaged the resulting structures by cryo-electron microscopy (cryo-EM), which confirmed that the elongated structures were composed of tightly packed microtubule bundles in the elongated structures (Figure 2D). We were able to 3D reconstruct three single microtubules, but due to tight packing of MTs within a bundle it was difficult to definitively select a single MT for 3D reconstruction. Those three single MTs generated by the CPC were composed of 14 or 15 protofilaments (Figure 2E). Polymerized MTs formed bundles, no astral-like MTs were observed in any of EM images. The MTs bundles were coated with unstructured material. Based on fluorescence microscopy images showing colocalization of MTs and CPC [Figure 2B], we posit this material to be the CPC. Taken together these results show that phase separation of CPC concentrates free tubulin and drive MT polymerization and bundling *in vitro*.

**Figure 2.**
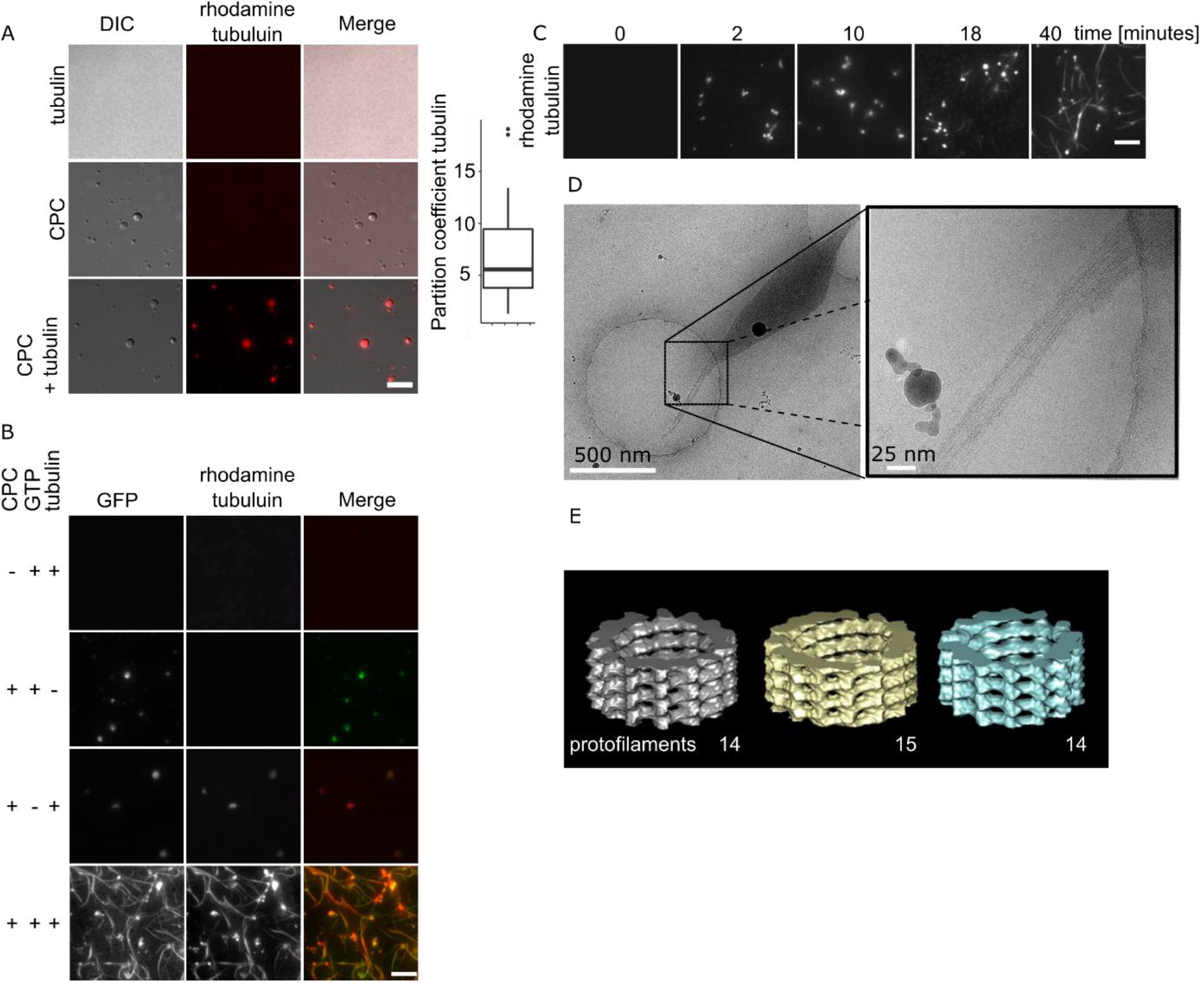
CPC coacervates generate bundles of MTs. A. DIC and fluorescence images of tubulin dimers partitioning into CPC coacervates (n=43). Experiment was repeated multiple times during the course of the project using different CPC preps (n=3 preps). B. DIC and fluorescence images of GFP partitioning into CPC coacervates (n=69). Experiment was repeated twice. C. Bundles of MTs generated in the presence of phase-separated CPC imaged by TIRF microscopy. Experiment was repeated multiple times during the course of the project using three different CPC preps; scale bar for A, B, C = 5 μm. D. Cryo-EM images of bundles of MTs generated by CPC. E. 3D reconstruction of single MTs generated by the CPC. Cryo-EM measurements were done twice from two independent CPC preps.

### MTs promote CPC liquid-liquid phase separation

α-satellite DNA, chromatin, or microtubule bundles induce phase separation of a CEN domain of the CPC. Similarly, MTs enhance the formation of condensates of TPX2 [30]. We investigated whether MTs can influence the CPC condensation state by mixing GFP-CPC at a concentration that is too low to phase separate on its own with GMpCpp stabilized microtubules and visualized the formation of condensates by fluorescence and DIC microscopy. CPC condensates were detectable on or near MTs. Under these conditions, the preformed MTs bundled together and were coated with CPC similar to the bundles that were formed by the CPC with free tubulin and GTP (Figure 3). We conclude that microtubules promote phase separation of the CPC.

**Figure 3.**
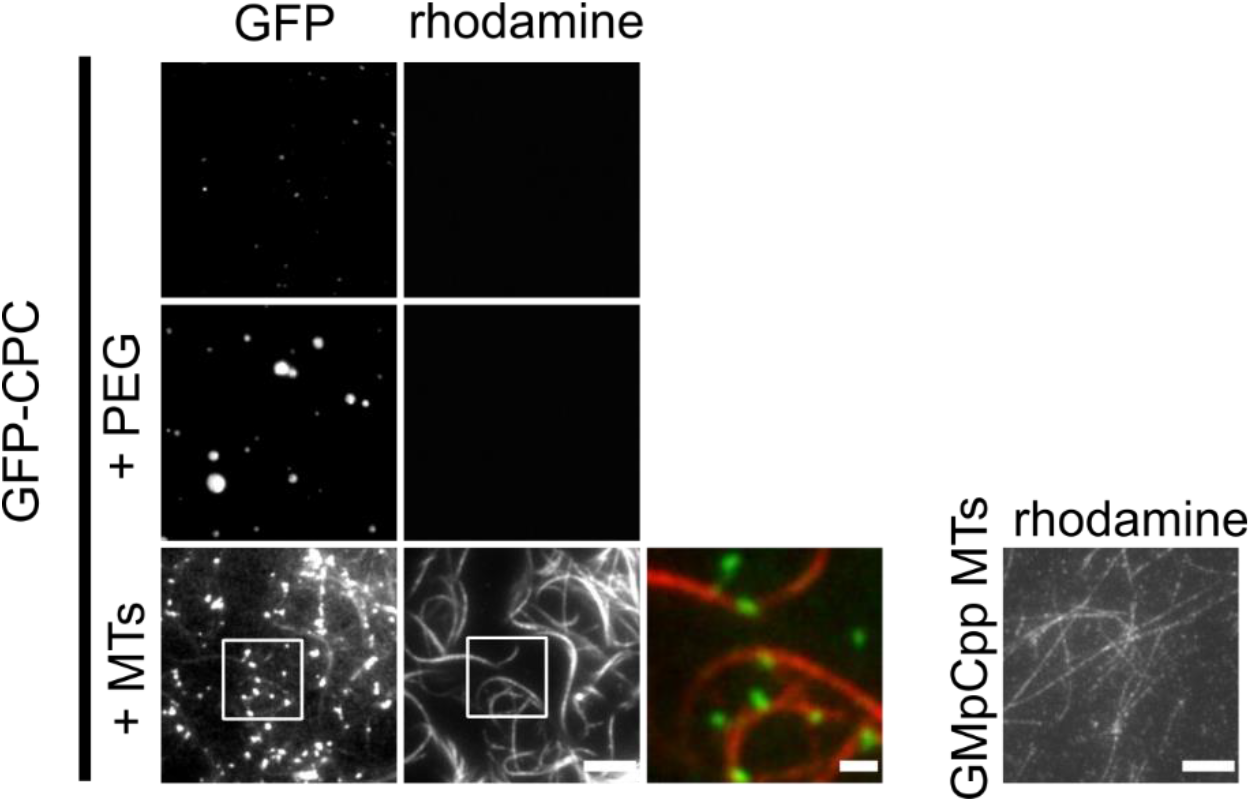
MTs induce phase separation of CPC. 100nM of GMpCpp stabilized MTs induced phase separation of 1 μM soluble CPC as imaged by TIRF microscopy with excitation at 480 nm (GFP-CPC) and 555 nm (rhodamine tubulin); scale bar for grey scale images = 5μm; scale bar for pseudo colored inset = 1 μm. Image to the right shows GMpCpp stabilized rhodamine labelled single MTs; scale bar = 5μm. Experiment was repeated twice.

### Polarity of bundles of MTs generated by the CPC

To understand the polarity of MTs protruding from CPC coacervates, we added purified kinesin-1-GFP (Supplementary Figure 4A), a plus-end directed motor, to MT bundles generated by the CPC and followed the direction of kinesin-1 movements by TIRF microscopy (Figure 4A, B), a technique known as motor-PAINT [31]. We compared the direction of movements on a single bundle to determine if all of the microtubules had a similar orientation. We grouped bundles based on direction of kinesin-1 movements into three classes; toward or away from CPC condensate, or multidirectional. Majority of the bundles had all (72%) or most (11%, defined as >=75% of movements in one direction) kinesin-1 movements oriented towards the CPC condensate (Figure 4C), suggesting that the plus-ends of polymerized MTs are inside or on the surface of CPC coacervates, and that the bundles are composed of parallel microtubules. Bundles showing kinesin-1 movements that were predominantly away from the condensates or multidirectional constituted 6% and 11% of bundles respectively (Figure 4C); however due to experimental setup the away CPC movement classification is more likely a result of misassignment of the originating condensate (Supplementary Figure 4B). These results suggest that MTs generated by CPC coacervates are parallel bundles oriented with plus-ends adjacent to or inside the nucleating condensate.

**Figure 4.**
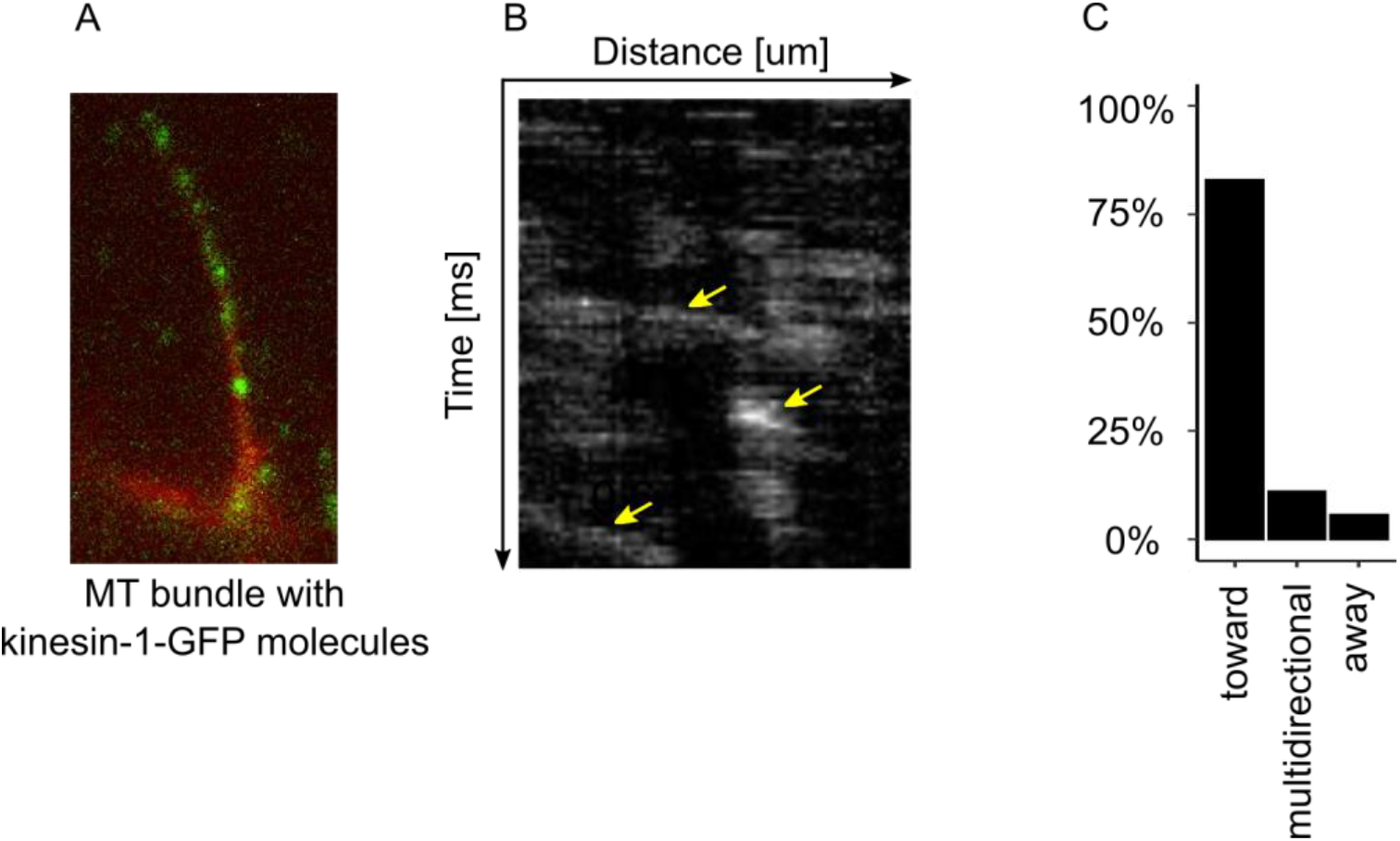
MTs generated by LLPS CPC are oriented with plus-ends inside the CPC condensate. A. An example of single Cy5-labelled MT bundle with kinesin-GFP molecules (green dots). B. Representative kymograph (from two independent experiments with 92 tracks) of single molecules of kinesin-1-GFP moving on bundle of MTs generated by the CPC. Yellow arrows represent spatial position of kinesin-1-GFP molecules on MT bundles over time C. Classification of MTs bundles based on predominant direction of kinesin-1-GFP movements relative to CPC condensate: toward (defined as >=75% of recorded kinesin movements on given bundle toward CPC condensate) or away (>=75% movements away from CPC condensate), or in both directions (multidirectional, remaining bundles).

### Borealin’s phase separation and MT binding activities are required to assemble MT bundles

We purified a set of mutant CPC CEN subcomplexes to investigate the mechanism by which the CPC generates parallel bundles of microtubules (Figure 5A, Supplementary Figure 5).

**Figure 5.**
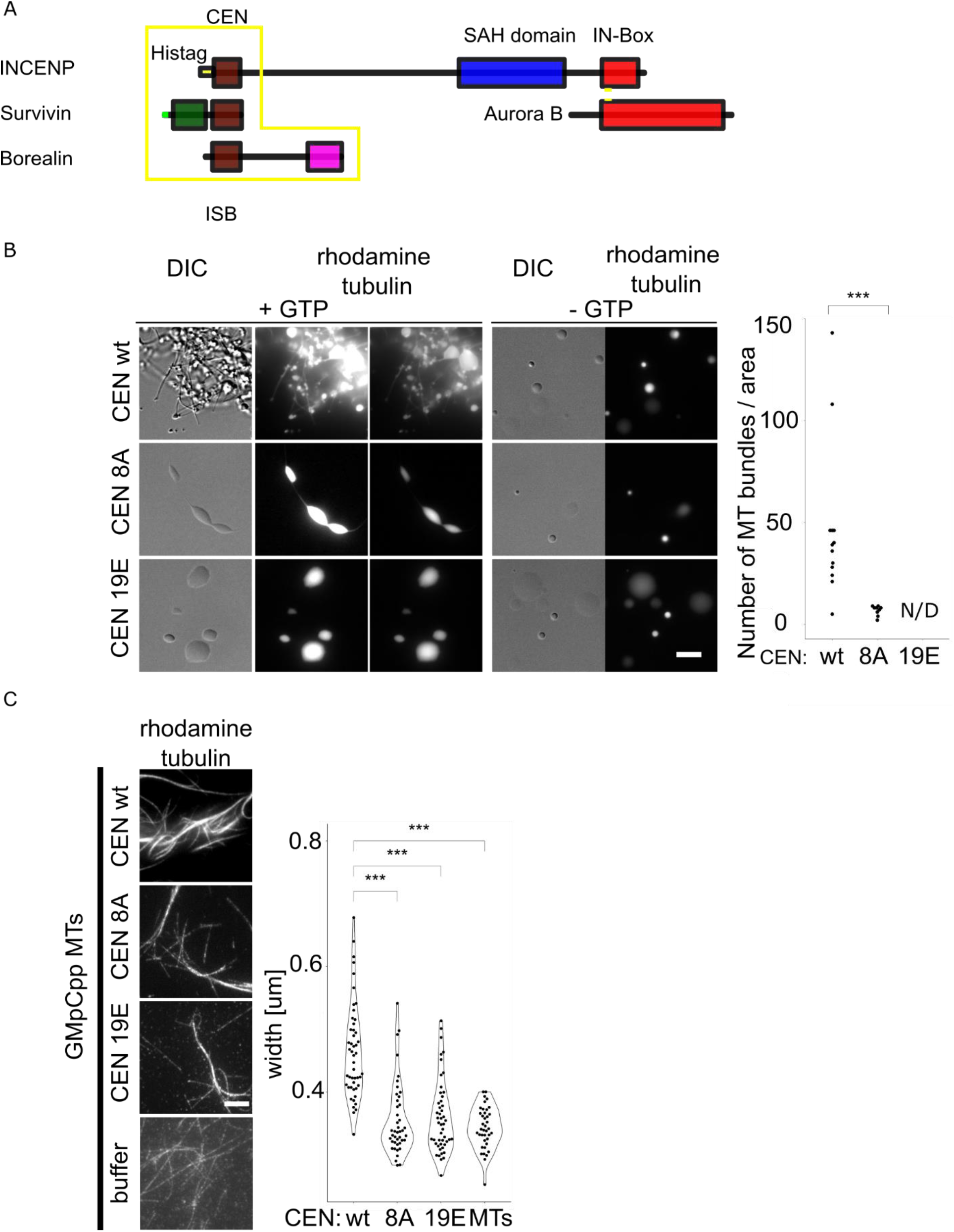
Phase separation propensity and MT binding of Borealin subunit of the CPC are important for MT nucleation and bundling. A. Scheme of the CPC; CEN: centromere-targeting domain composed of full length Survivin and Borealin, N-terminal fragment of INCENP. B. DIC and fluorescence images of rhodamine labelled MTs generated by CEN-Borealin^wt^, CEN-Borealin^8A^ and CEN-Borealin^R17E,R19E,R20E^. Experiment was repeated three times. Plot shows the number of MTs bundles per given area; Total number of MT bundles for CEN-Borealin^wt^ n = 612, for CEN-Borealin^8A^ n = 61; p-value^wt-8A^ = 5.5·10^-5^. C. GMpCpp stabilized rhodamine labelled MT bundled by CEN-Borealin^wt^, CEN-Borealin^8A^ and CEN-Borealin^R17E,R19E,R20E^ imaged in TIRF mode with excitation at 555 nm (rhodamine tubulin). Plot shows quantification of the width of single bundles of MTs. Experiment was repeated twice; CEN-Borealin^wt^ n = 49, CEN-Borealin^8A^ n = 43, CEN-Borealin^R17E,R19E,R20E^ n = 49, MTs n = 39; p-value^wt-8A^ = 1.05·10^-9^; p-value^wt-19E^ = 1.68·10^-9^. Statistical analysis was performed by applying Kolmogorov-Smirnov test; * = p<0.05, ** = p < 0.01, *** = p<0.001.

To determine whether Borealin phase separation was important for MT nucleation and bundling we employed a CEN mutant where 8 lysines in 21 amino acids stretch of an intrinsically disordered region of the Borealin subunit are mutated to alanines that is deficient in LLPS(CEN-Borealin^8A^)[18]. CEN-Borealin^8A^ generated significantly less MTs than CEN-Borealin^wt^ (Figure 5B). Next, we checked the role of the MT-binding region of Borealin in the formation of MT bundles. We used charge-reversed CEN-Borealin^R17E,R19E,R20E^ mutant that has reduced MT binding affinity [16]. This mutant was able to phase-separate similar to the CEN-Borealin^wt^ but no MTs were detected in the *in vitro* polymerization assay (Figure 5B).

To investigate the role of Borealin phase separation and MT binding in formation of MT bundles independently of MT nucleation process, we incubated CEN-Borealin^wt^ and mutants with GMpCpp-stabilized MTs and quantified the width of MT bundles. CEN-Borealin^wt^ generated on average bundles of 0.4 μm width, while bundles generated by mutants were significantly thinner. The measured width of single GMpCpp is 0.2 μm, which is the resolution limit of the fluorescence microscope (Figure 5C). These results suggest that the Borealin subunit of the CPC contributes to the formation of MT bundles *in vitro* through a combination of its abilities to phase separate and to bind MTs.

### Borealin-dependent liquid-liquid phase separation and MT binding regulates midzone microtubules independent of Aurora B kinase activity

We employed the phase separation and microtubule binding mutants to test the importance of the CPC’s microtubule bundling activities in HeLa cells. Cells arrested in S-phase were depleted of endogenous Borealin by siRNA accompanied by the expression of siRNA-resistant recombinant wild-type or mutant Borealin. The cells were released from the block, fixed in glutaraldehyde to preserve MT structure, and analyzed during the first mitotic transition (Figure 6A, Supplementary Figure 6A). Previous characterization of these mutants has shown that the localization of the CPC is reduced by these mutations at inner-centromeres and midzones [18, 19]. Spindle structure appeared to be unaffected, however, the midzones of cells mutated for MT binding or phase separation contained approximately two-fold less tubulin than those of cells rescued with Borealin^wt^ (Figure 6B). The amount of tubulin in the midzone was quantified relative to total tubulin in the anaphase spindles. These changes in the midzone were not an indirect effect of changes to Aurora B activity since adding an Aurora B inhibitor (AZD1152-HQPA-Barasertib) did not affect the amount of midzone microtubules (Supplementary Figure 6B). These data demonstrate that the MT population of anaphase midzones is dependent on both the microtubule binding and phase separation activities of the CPC, consistent with a requirement for CPC phase separation-dependent MT bundling activity.

**Figure 6.**
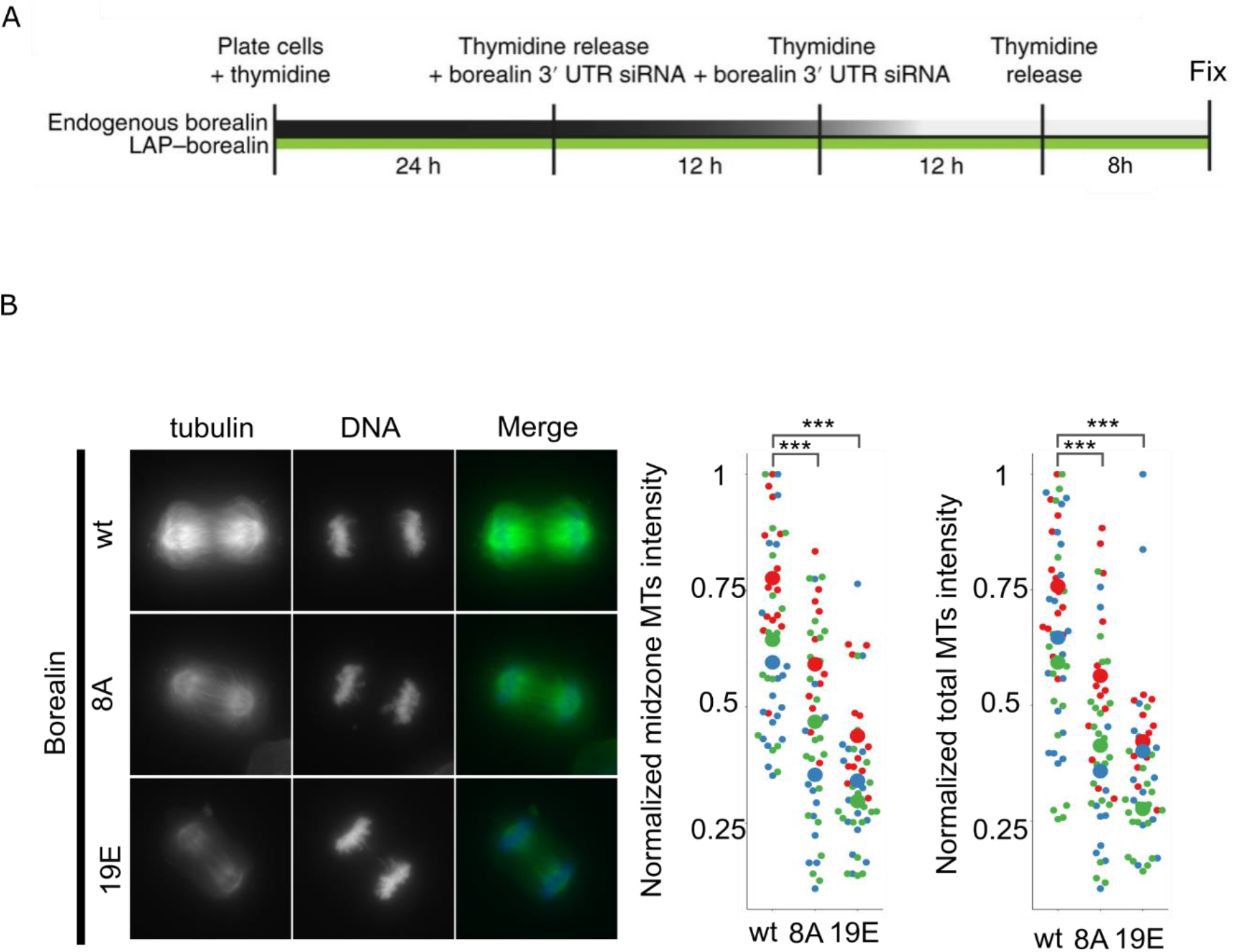
CPC phase separation and MT bundling are important for formation of proper midzone during anaphase and k-fibers in nocodazole washout cells. A. Experimental design for knockdown and replacement of Borealin by siRNA in HeLa cells. B. Fluorescence images of HeLa cells with endogenous Borealin replaced with wild type or indicated Borealin mutants and stained for α-tubulin and DAPI. Experiment was repeated three times; CEN-Borealin^wt^ n=(28, 38, 32); CEN-Borealin^8A^ n=(26, 28, 42); CEN-Borealin^R17E,R19E,R20E^ n=(26, 30, 38). p-value^wt-8A^ = (0.015, 0.00022, 0.0058); p-value^R17E,R19E,R20E^ = (1.77·10^-6^, 6.9·10^-5^, 3.3·10^-8^). For statistical analysis Welch’s t-test with Bonferroni correction was applied; * = p<0.05, ** = p < 0.01, *** = p<0.001.

Another parallel MT binding structure of the mitotic spindle is kinetochore-associated MTs which are observed after washing cells out of nocodazole. Both Borealin mutants were also deficient in formation of kinetochore-associated MTs (Supplementary Figure 6C) demonstrating that the CPC’s microtubule organizing activity is not specific to anaphase midzones.

## Discussion

We demonstrated that the CPC employs its ability to bind MTs and undergo LLPS in order to organize parallel bundled MT structures that are central features of the anaphase midzone. A simple *in vitro* system of phase separated CPC, tubulin, and GTP generated robust tapered parallel bundles of MTs that elongated from condensates over time. We showed that CPC phase separation driven by Borealin is important for maintaining proper density of MTs in midzone during anaphase and for generation of kinetochore-associated MTs that form on chromosomes after removal of nocodazole. In broader context our results extend the emerging concept that LLPS of proteins provides a mechanism to concentrate molecules to regulate cellular process, and to act as organizational hubs for formation of ordered cellular structures [32].

In our experimental setup, MT nucleation and bundling were dependent on CPC droplets and specific interactions between the CPC and tubulin, as soluble CPC did not nucleate or bundle MTs. Indeed, it has been shown that there are two MT-binding regions within CPC: one on the N-terminal fragment of Borealin [16], and another within the INCENP SAH domain [17]. Using a CEN domain construct with MT-binding deficient Borealin, we demonstrated that interactions between tubulin and Borealin are important to initiate MTs nucleation and bundling. Current models assume that tubulin dimers have to assembly into longitudinal oligomers that subsequently form a sheet through lateral association with tubulin dimers [33]. We suggest that the CPC stabilizes the generated MTs through its bundling activity. Another possibility is that by phase separation CPC creates a space so dense in tubulin binding motifs that tubulin dimers interact with each other to form microtubule seeds, and to begin nucleation *in vitro*. Currently, we have no evidence that CPC-dependent MT nucleation happens *in vivo* and we currently favor models where the CPC works with MT nucleators such as γ-TuRC. We noticed that the protofilament numbers of MTs generated by the CPC *in vitro* are different from those generated by the γ-TuRC complex *in vivo*, which suggest that MTs bundling, not nucleation, is an important function of CPC in regulation of MT bundles. We suggest that the CPC acts similarly to the TPX2 protein that also phase separates and nucleates MTs *in vitro* and it has been shown to work with the augmin complex to recruit γ-TuRC to nucleate branched MT structures *in vivo*. We suggest that the CPC works with regulatory proteins such as γ-TuRC to control the number of protofilaments, and a function of phase separated CPC is to sequester free tubulin and bundle MTs within midzones. We suggest that there are important similarities between the complexes that form parallel and branched MT structures in the spindle. We suggest that the CPC combines the activities of augmin and TPX2, and acts as both a parallel microtubule bundler (rather than augmin’s side binding), and a complex that phase separates to concentrate free tubulin dimers (analogous to TPX2). An important area of future research will be to test if the γ-TuRC also works in combination with the CPC [34].

We previously showed that LLPS of CPC is important for its mitotic functions such as correcting kinetochore–microtubule attachments and maintaining the spindle checkpoint [18]. Here we show that the CPC combines phase separation and its ability to interact with tubulin to form parallel MT bundles. Our *in cell* analysis revealed that Borealin-dependent phase separation and MT binding are important for formation of midzone and kinetochore-associated MTs in HeLa cells. Our studies showed that MT bundles generated by the CPC *in vitro* are oriented such that plus ends are inside coacervates and minus ends are protruding out of condensates (Figure 4). In dividing cells, microtubule bundles formed at kinetochores (so called pre-formed k-fibers) have been observed in monastrol-treated cells, as well as in untreated cells [35]. Those pre-formed k-fibers were incorporated into the spindle by a NuMA-dependent mechanism [35]. Such mechanism has been shown to constitute centrosome-independent pathway that promotes MT nucleation, and to work as self-organizing center that assists chromosomes biorientation [35, 36]. The ability of phase-separated CPC to nucleate and bundle parallel MTs with minus ends protruding from droplets, centromeric localization of the CPC during prometaphase/metaphase [36, 37], and defects of kinetochore associated MTs in cells expressing mutants of either Borealin’s MT binding or its phase separation activities [27] suggests that CPC contributes to formation of pre-formed k-fibers.

In conclusion, we have uncovered the novel ability of the CPC to extend tapered microtubule bundles from a central organizing condensate. How the CPC is able to allow plus end growth within a condensate while “pushing” the bundles outwards is another important area of future research.

## Materials and methods

### Protein purification

Chromosomal passenger complex: *Xenopus* INCENP and Aurora B (amino acids 33-368) genes were cloned into pMCSG11 vector as a bi-cistronic construct with an N-terminal 6xHIS affinity tag on the INCENP protein. *Xenopus* Survivin-B and Borealin/Dasra-A were cloned into pET15b as a separate bi-cistronic vector. The recombinant CPC from *Xenopus laevis* was overexpressed in *Escherichia coli* strain BL21-CodonPlus (DE3) Magic containing both vectors in the presence of carbenicillin (100 μg/mL), kanamycin (50 μg/mL) and chloramphenicol (34 μg/mL). Bacteria were induced at OD 0.6 with 0.15 mM isopropyl-β-d-thiogalactoside followed by adding 80 μM ZnCl_2_ per 1 L of Luria-Broth (LB) medium. Cells were incubated with shaking for 16 h at 18°C. Cells were harvested, and pellets were suspended in lysis buffer (2 mM imidazole, 50 mM Tris, pH 7.9, 500 mM NaCl, 5% glycerol, 0.5 mM TCEP) containing Complete Inhibitor Protease Cocktail (Roche, Indianapolis, IN), lysozyme and DNAse. Cells were lysed using an EmulsiFlex-C3 homogenizer. Lysates were clarified by ultracentrifugation at 30,000 rpm for 40 min at 4°C. The supernatant was incubated with Ni^2+^ resin (Qiagen) at 4°C. Beads were washed in a buffer containing 10 mM imidazole 50 mM Tris, pH 7.9, 500 mM NaCl, 5% glycerol, 0.5 mM TCEP, and proteins were eluted with a buffer composed of 250 mM Imidazole 50 mM Tris, pH 7.9, 500 mM NaCl, 5% glycerol, 0.5 mM TCEP. Protein was separated on Superdex200 in a buffer 50 mM Tris, pH 7.9, 500 mM NaCl, 5% glycerol, 0.5 mM TCEP. Fractions containing CPC were pooled and concentrated centrifugal filter unit with cut-off 30 kDa (Amicon), aliquoted, and stored at −80.

Kinesin-1-GFP: Plasmid coding for kinesin-1-GFP was purchased from Addgene repository (#129761). The protein was expressed in BL21(DE3)RIL cells. Bacteria were induced at OD 0.6 with 0.15 mM isopropyl-β-d-thiogalactoside in Luria-Broth (LB) medium. Cells were incubated with shaking for 16 h at 18°C. Cells were harvested, and pellets were suspended in a lysis buffer (2 mM imidazole, 50 mM Tris, pH 7.9, 200 mM NaCl, 1% glycerol, 0.5 mM TCEP, 4 mM MgSO_4_) containing Complete Inhibitor Protease Cocktail (Roche, Indianapolis, IN), lysozyme and DNAse. Cells were lysed using an EmulsiFlex-C3 homogenizer. Lysates were clarified by ultracentrifugation at 35,000 rpm for 40 min at 4°C. The supernatant was incubated with Ni2+ resin at 4°C. Beads were washed in a wash buffer containing 10 mM imidazole 50 mM Tris, pH 7.9, 500 mM NaCl, 1% glycerol, 0.5 mM TCEP, 4 mM MgSO_4_, then washed again with a wash buffer supplemented with 50 μM ATP. Protein was eluted with a buffer composed of 80 mM PIPES pH=7.0, 4 mM MgSO_4_, 300 mM imidazole, 50 μM ATP. Protein was separated on Superdex200 in a buffer 80 mM PIPES pH=7.0, 4 mM MgSO_4_, 1 mM EGTA. Fractions containing kinesin-1-GFP were pooled and concentrated.

CEN-Borealin is composed of human INCENP (1-58 aa), full-length human Survivin and full-length human Borealin. CEN-Borealin^wt^, CEN-Borealin^8A^ and CEN-Borealin^19E^ were purified using the same protocol as for purification of the CPC. The CEN protein was expressed from pET28b vector in BL21(DE3) Rosetta cells.

### Phase separation assay

The CPC phase separation was induced by incubating 1 μM CPC in a buffer containing 150 mM NaCl and 20 mM HEPES 7.2, or a buffer composed of 500 mM NaCl and 20 mM HEPES 7.2 supplemented with 7% PEG3350 to serve as a crowding agent (Figure 1C). For consistency, in all other experiments CPC phase separation was induced by mixing with a buffer composed of BRB40 (40 mM PIPES, 0.5 mM MgCl_2_, 0.5 mM EGTA, pH 6.8), 1 mM GTP, 1mM DTT and 7% PEG3350. For time-lapse imaging of CPC droplet fusion, CPC phase separation was induced with a BRB40 mix containing 7% PEG3350, the mixture was transferred to microscopic chambers and droplet fusion was immediately imaged under x63 objective (Plan Apo) DIC on a ZEISS AxioObserver Z1 microscope.

### MTs polymerization assay

1 μM CPC in SEC buffer was mixed with BRB40 buffer (40 mM PIPES, 1 mM MgCl2, 1 mM EGTA, pH 6.8 with NaOH), 1 mM GTP, 1mM DTT and 7% PEG3350 to induce phase separation. Then 2 μM of α/β tubulin dimers (PurSolutions) in BRB80/DTT buffer was added and reaction was kept at RT for 40 minutes. Unlabeled tubulin was mixed with labelled tubulin at 10:1 ratio. Polymerized MTs were transferred onto cover glasses or chambers (Grace Bio-Labs) and observed using a ×63 objective on a Zeiss Observer Z1 wide-field microscope by fluorescence and DIC imaging or using a x100 objective on a LEICA Thunder/TIRF.

For microtubule polymerization by CEN complex, CEN^wt^, CEN^8A^, CEN^19E^ at 7 μM concentration were incubated with BRB40, 1 mM GTP, 1mM DTT, 7% PEG3350 and tubulin for 40 minutes at RT. Reaction was transferred in between cover glasses and imaged on x63 objective on a ZEISS microscope.

### Kinesin-1 motility assay – motorPAINT assay

Cy5-labelled MTs were polymerized in the presence of GTP and phase-separated CPC. Reaction mixture was transferred into coverslip-bottomed CultureWell (Grace BioLabs), then solution of glucose/oxidase system (40 mM glucose, 130 mg/mL glucose oxidase and 24 mg/mL catalase), 1 mM ATP and 30 nM of kinesin-1-GFP were added. Kinesin movements were imaged using x100 objective in a TIRF mode with excitation at 480 nm on the LEICA Thunder microscope and analyzed in FIJI.

### Partition coefficient of tubulin and peptides into CPC coacervates

Partition coefficient for H3T3ph(1-21)-FITC, H3T3(1-21)-FITC and rhodamine tubulin into CPC condensates was measured according to the protocol described in Trivedi et al. 400 nM of peptide, or 2 μM of α/β tubulin dimers (1:10 rhodamine labeled to unlabeled) in BRB80/DTT was added to CPC condensates that were generated by mixing 1 μM of the CPC with BRB40 buffer, 1mM DTT and 7% PEG3350. Condensates were imaged on x63 objective on a ZEISS microscope Fluorescence signals were calculated in FIJI. Partition coefficients were calculated by dividing the fluorescence signal per unit area inside the coacervates by the fluorescence signal per unit area outside the coacervates after subtracting the background fluorescence. Background fluorescence was calculated by imaging the coacervates in the absence of fluorescent agent molecules.

### Bundling of MTs by CEN subcomplex

100 nM of GMpCpp stabilized, rhodamine labelled MTs were mixed with 4 μM CEN^wt^, CEN^8A^, CEN^19E^ which is below critical concentration for CEN. After 2 minutes incubation reactions were transferred in between cover slides and imaged on the x100 objective using TIRF mode on LEICA Thunder microscope. Bundling reactions were performed in BRB80/DTT buffer. GMpCpp MTs were prepared by incubating 200 μM α/β tubulin dimers with 1 mM of GMpCpp for 1h at 37C in the BRB80/DTT buffer.

### Kinase assay and immunoblotting

1 μM of recombinant CPC was mixed with 2 μg of GST-histone H3 (1-21aa) fragment, 10 mM ATP, 10 mM MgSO_4_, 2 mM DTT and 20 mM HEPES 7.2 and 7% PEG3350 and incubated at 37C for 5 minutes. The reaction mixture was resolved on 15% SDS-PAGE gel and blotted with antibodies against histone H3 S10ph (Cell Signaling Technologies, 1:1000).

### EM data collection and MTs reconstruction

The polymerized by CPC MTs sample (~3.0 μL) was applied to discharged lacey carbon grids and plunge frozen in liquid propane. Frozen grids were imaged in a Titan Krios at 300 keV. MTs reconstruction was done in ImageJ using a TubuleJ plugin according [38, 39].

### CPC mass spectrometry analysis

xCPC protein sample was prepared as two 5 μg aliquots for trypsin and GluC peptide mapping as previously described [40]. Samples were analyzed by nanoLC-MS/MS (nanoRSLC, ThermoFisher) coupled to an Orbitrap Eclipse using a 120 min reversed phase gradient with resolution settings of 120,000 and 15,000 (at m/z 200) for MS1 and MS2 scans, respectively. Selected peptides were fragmented using stepped high energy collisional dissociation (20, 30, 40%). Tandem mass spectra were analyzed according to a label-free proteomic strategy using Proteome Discoverer (version 2.5.0.400, ThermoFisher) with the Byonic (version 4.1.10, Protein Metrics) and Minora nodes and using a database consisting of E. coli and Xenopus CPC protein sequences to objectively identify all CPC and background E. coli proteins present in the purified sample [41, 42]. Mass tolerances of 10 ppm and 20 ppm were used for matching parent and fragment masses, respectively. Spectra were searched with a fixed modification of carbamidomethyl (C) and variable modifications of phosphorylation (S, T, Y). Tandem mass spectra were then analyzed according to a peptide mapping strategy. MS/MS spectra were extracted and charge state deconvoluted, then searched with a database consisting of the top 20 Byonic-identified proteins (sorted by Log Probability and accounting for >99% of protein signal) using the MassAnalyzer algorithm embedded in BioPharma Finder software (version 4.1.53.14, Thermo Fisher) [43, 44]. Searches were performed with a MS Noise Level of 2000 and a S/N threshold of 10, parent ion tolerance of 10 ppm and an MS/MS Minimum Confidence Level of 80%. Trypsin was selected as the protease with a ‘High’ specificity setting. Variable modifications of carbamidomethyl (C), oxidation (M), deamidation (N, Q), and phosphorylation (S,T,Y) was specified. Phosphopeptide identifications were manually validated.

### Cell culture

HeLa T-REx cells (ThermoFisher Scientific) stably expressing LAP-Borealin^wt^, LAP-Borealin^8A^ and LAP-Borealin^19E^ [16] were grown in Dulbecco’s modified Eagle’s medium (DMEM; Invitrogen) supplemented with 10% fetal bovine serum (Gibco) in the presence of 5% CO2 in a humidified incubator at 37 °C.

### siRNA transfection

For knockdown and replacement experiments, Borealin 3’ UTR siRNA (AGGUAGAGCUGUCUGUUCAdTdT) was transfected using RNAiMAX (Invitrogen) according to the manufacturer’s protocol. Mitotic phenotypes were analyzed in the first mitosis after complementation. Borealin stable cell lines were plated in the presence of 2 mM thymidine for 24 hours. Next, the fresh DMEM/10% FBS media was added and cells were transfected with siRNA, and incubated for 12 hours. Next, media was replaced and cells were incubated with DMEM/10% FBS with 2 mM thymidine, transfected with siRNA and incubated for 12 hours. Next, cells were released from thymidine in fresh media DMEM/10% FBS for 9 hours and fixed. For nocodazole washout experiments cells were incubated with 3.3 μM nocodazole for 5 minutes. Nocodazole was washed away 3 times with DMEM/10% FBS and fixed. For Aurora B inhibition experiments, 1 μM AZD1152-HQPA (Barasertib, SelleckChem) was added for 30 minutes before fixation in glutaraldehyde.

### Immunofluorescence

Cells were plated on the cover slides coated with poly-L-lysine (Sigma). For midzone staining and analysis cells were extracted for 30 seconds with 0.5% Triton X-100 in PHEM buffer (25 mM HEPES, 60 mM PIPES, 10 mM EGTA and 4 mM MgCl2, pH 6.9) and then fixed in **2.5%** glutaraldehyde in PHEM buffer for 10 minutes. Reaction was quenched with sodium borohydride for 7 minutes. Slides were washed 3 times with TBS(T) buffer, blocked for 1 hour in 3% BSA/TBS(T) buffer, incubated with anti-tubulin DM1α-FITC labelled antibody for 1 hour. Slides were washed 4 times with TBS(T) buffer, the last wash contained DAPI (0.5 μg/mL). Next cells were washed twice with water, and mounted with ProLong™ Gold Antifade (Thermo Fisher Scientific). For pre-formed k-fibers staining cells were fixed in 4% PFA in PHEM buffer with 0.5% Triton X-100 for 20 minutes at RT. Slides were blocked for 1 hour in 3% BSA/TBS(T) buffer, incubated with primary antibody: rabbit anti-α-tubulin DM1α, mouse anti-γ-tubulin, and human anti-ACA for 1 hour. Slides were washed 5 times with TBS(T) buffer, and incubated with secondary antibodies for 45 minutes. Cells were washed 4 times, the last wash contained DAPI (0.5 μg/mL). Next cells were washed twice with water, and mounted with ProLong™ Gold Antifade.

### Image analysis

Images were acquired with a 63X/1.4 NA objective on a Zeiss Observer Z1 wide-field microscope. Image analysis was done in FIJI. The Z-stacks were projected using maximum intensity algorithm. **Midzone analysis:** To analyze the level of MTs in midzone of anaphase cells the intensity of MTs-FITC signal at the region between separating chromatids was divided by the intensity of total area of anaphase cell measured for FITC channel. **K—fibers analysis:** To analyze the total intensity of MTs polymerized at centromeres, the polymerization sites were localized based on 650 nm channel (corresponding to ACA staining) with pixel intensities higher than background. The background was subtracted using the rolling ball algorithm with the radius of curvature of the paraboloid bigger than the centromere size. Centrosomes were excluded from analysis based on gamma-tubulin staining. Centromere intensities were selected based on certain threshold values. **motor-PAINT:** Kinesin-1 movements along polymerized MTs were analyzed in ImageJ using multi kymograph plugin.

### Statistics and reproducibility

All of the key experiments were repeated multiple times as indicated in the figure legends. Experiments for Figure 1e and 1f, Figure 2b and 2d, Figure 3 and Figure 4, and Figure 5c were performed twice independently with similar results. Experiments for Figure 5b and Figure 6b were performed three times independently with similar results. Indicated statistical analysis was performed using R (v.3.6.3).

## Author contribution

EN and PTS designed the study. EN performed protein purification and all *in vitro* and in cell experiments and analyses presented in the study except for microtubule 3D reconstruction. TT purified CPC and GFP-CPC. LE prepared sample for mass spectrometry analysis. SR performed tomography data collection. DC performed 3D reconstruction of microtubules. EN, TT, LE and PTS wrote the manuscript. All authors read and approved the final version of the manuscript.

## Acknowledgements

We would like to thank Dr. Aaron Bailey for doing MS analysis. We would like to thank Dr. Dan Burke for helpful comments and discussions. EN, TT and PTS were supported by NIH R01GM139787. EN and PTS had additional support from a grant from the Schiff foundation. DC was supported by a French National Research Agency grant (ANR-18-CE13-0001-01). The Mass Spectrometry Facility is supported in part by Cancer Prevention Research Institute of Texas (CPRIT) grant number RP190682. Transmission electron micrographs were recorded at the University of Virginia Molecular Electron Microscopy Core facility (RRID: SCR_019031), which is supported in part by the School of Medicine. The Titan Krios (SIG S10-RR025067), Falcon II/3EC direct detector (SIG S10-OD018149), and K3/GIF (U24-GM116790) were purchased in part or in full with the designated NIH grants.

---

**Supplementary Figure 1.**
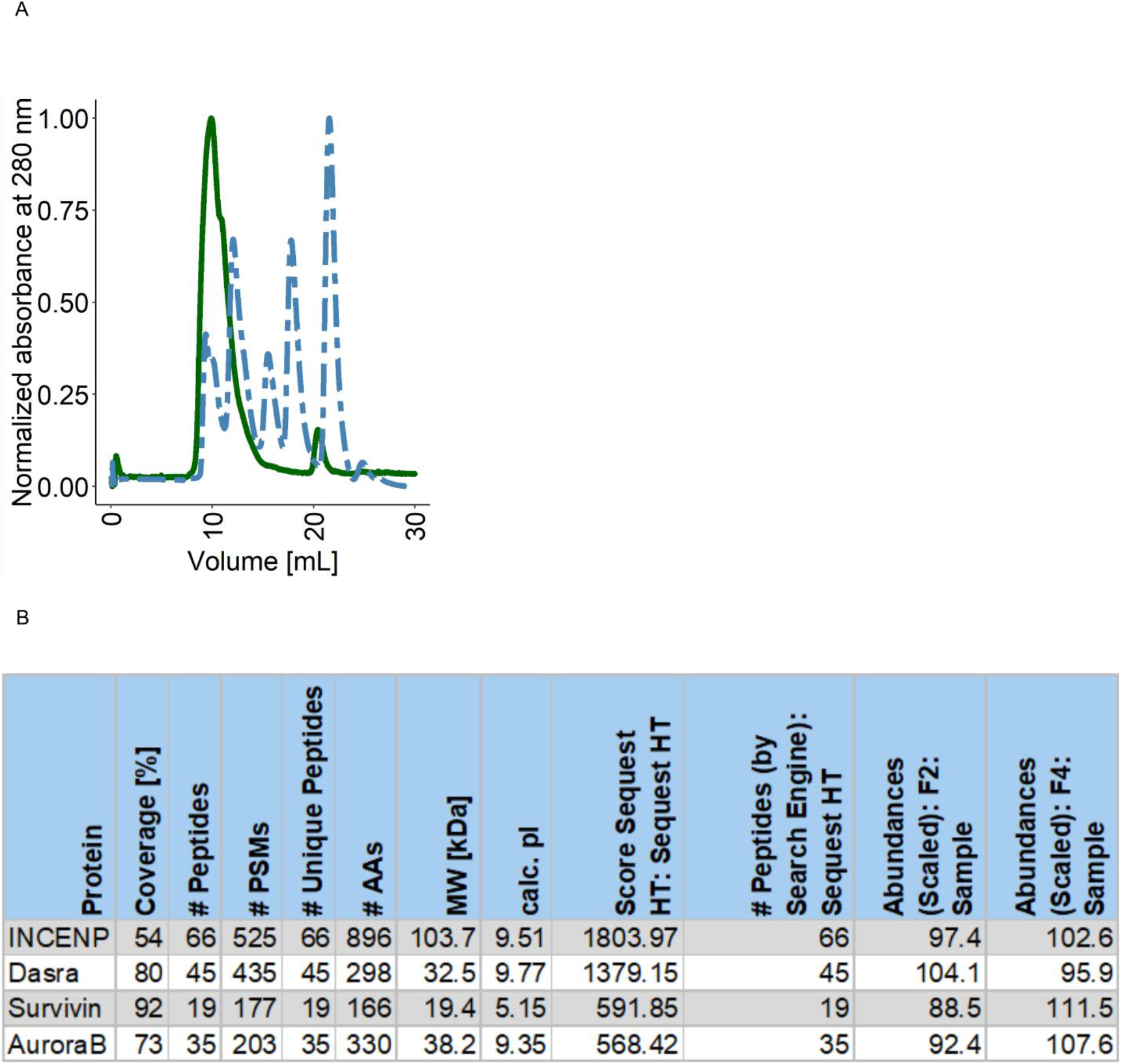
Biochemical characterization of recombinant CPC prep. A. Size exclusion profile of the CPC. Protein complex was separated on Superdex 200 10/300 GL attached to BioLogic DuoFlow™ Medium-Pressure Chromatography System (Bio-Rad) in a buffer composed of 20 mM HEPES 7.2, 500 mM NaCl, 2% glycerol, 0.5 mM TCEP. CPC elution profile is marked as solid line; elution profile of protein standards is marked as dotted line. B. Mass spectrometry results of the CPC protein prep.

**Supplementary Figure 2.**
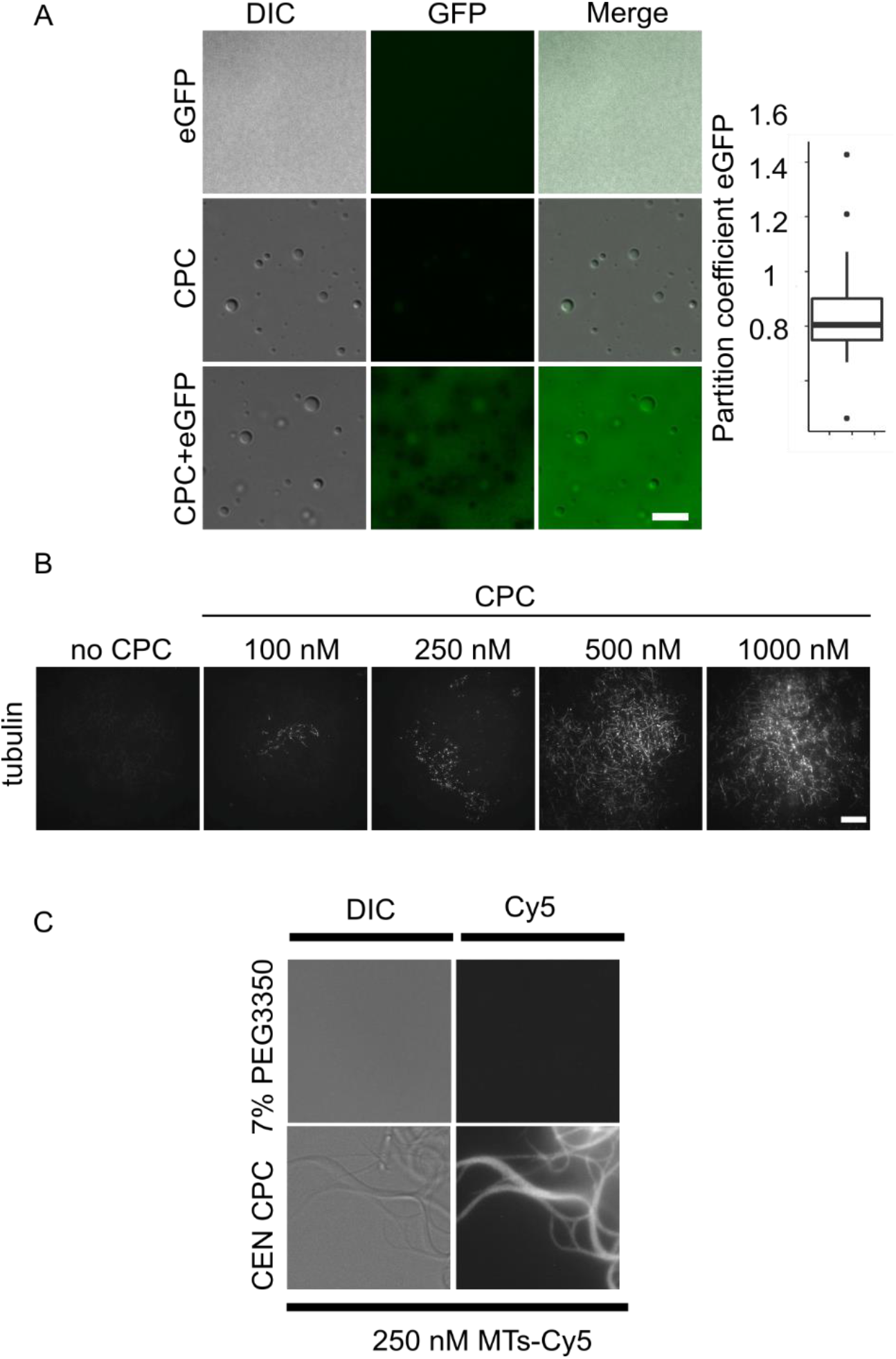
MTs are polymerized and bundled *in vitro* by the phase-separated CPC. A. DIC and fluorescence images showing partitioning of GFP into CPC condensates; objective 63x; scale bar = 5μm. B. Rhodamine labelled α/β-tubulin dimers were incubated with various concentrations of LLPS CPC, and resulting MT bundles were imaged by TIRF microscopy; objective 100x; scale bar 20 μm. Images present the largest observed structure from each concentration. Experiment was repeated twice and for each repetition 4 to 8 fields of view were imaged. C. DIC and fluorescence images of GMpCpp stabilized MTs labelled with Cy5 incubated with 7% PEG3350 or 10 μM CEN subcomplex; objective 63x; scale bar = 5μm.

**Supplementary Figure 4.**
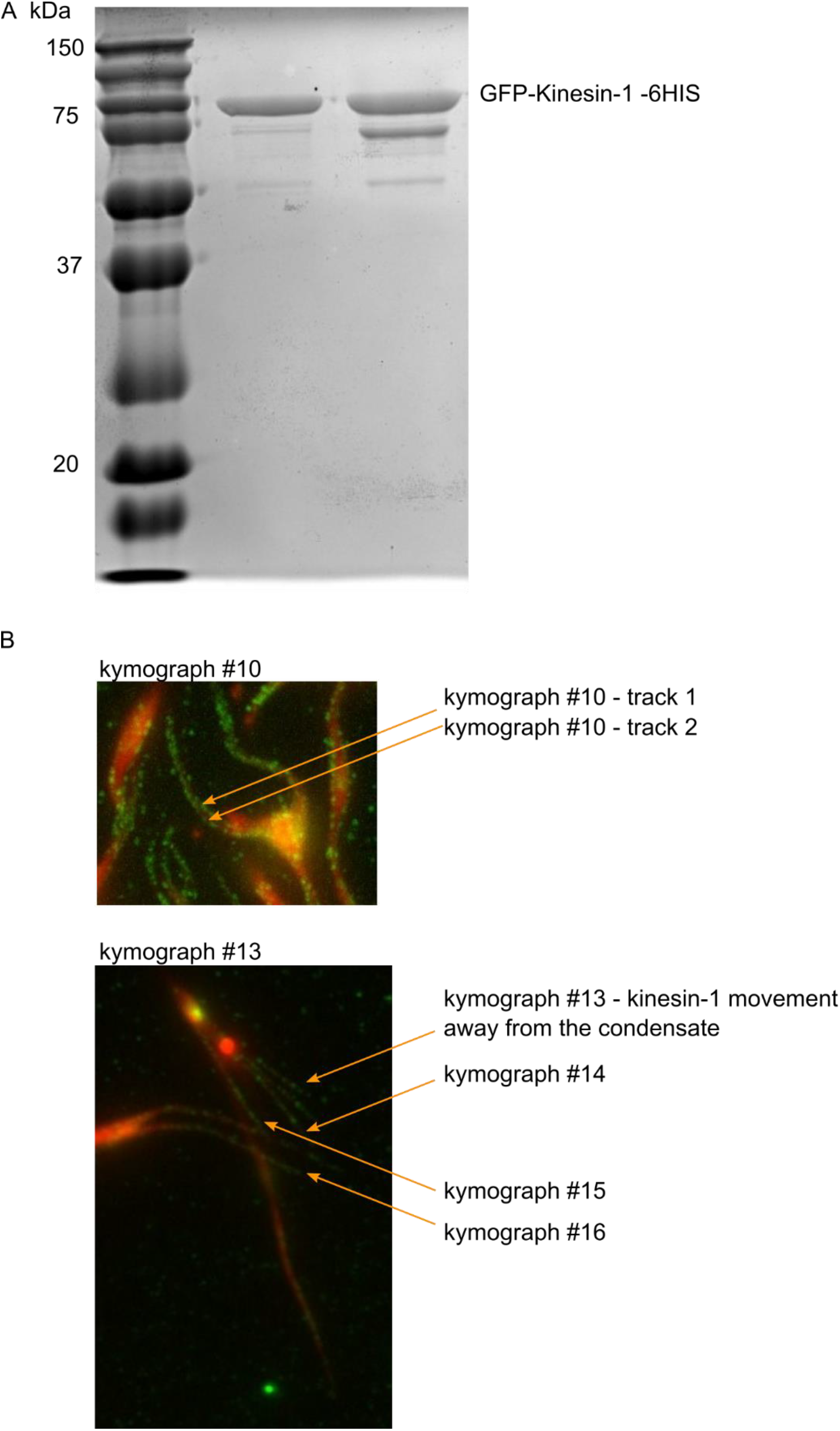
Kinesin-1-GFP purity and motorPAINT analysis. A. SDS-PAGE gel stained with Coomassie to show the composition and purity of the kinesin-1-GFP prep used in motor-PAINT experiment. B. Example of the large bundle (kymograph #10.) that had two separable kinesin-1 tracks with opposite directions (toward and away from the condensate). We believe that these types of large MT bundles are formed by fusing already formed MT bundles of same orientation of MTs. Due to nature of TIRF, we may not observe the remaining part of the bundle that is above the TIRF acquisition plane. Different MT bundles may still be fused together in orientation that results in the overlapping kinesin tracks thus resulting in bundles classified as multidirectional. The CPC condensate origin of the bundle may potentially be misidentified and bundle classified with away orientation (the bundle origin of kymograph #13 is less clear than the #14 - #16, and it may come from the condensate that is not visible in TIRF acquisition plane).

**Supplementary Figure 5.**
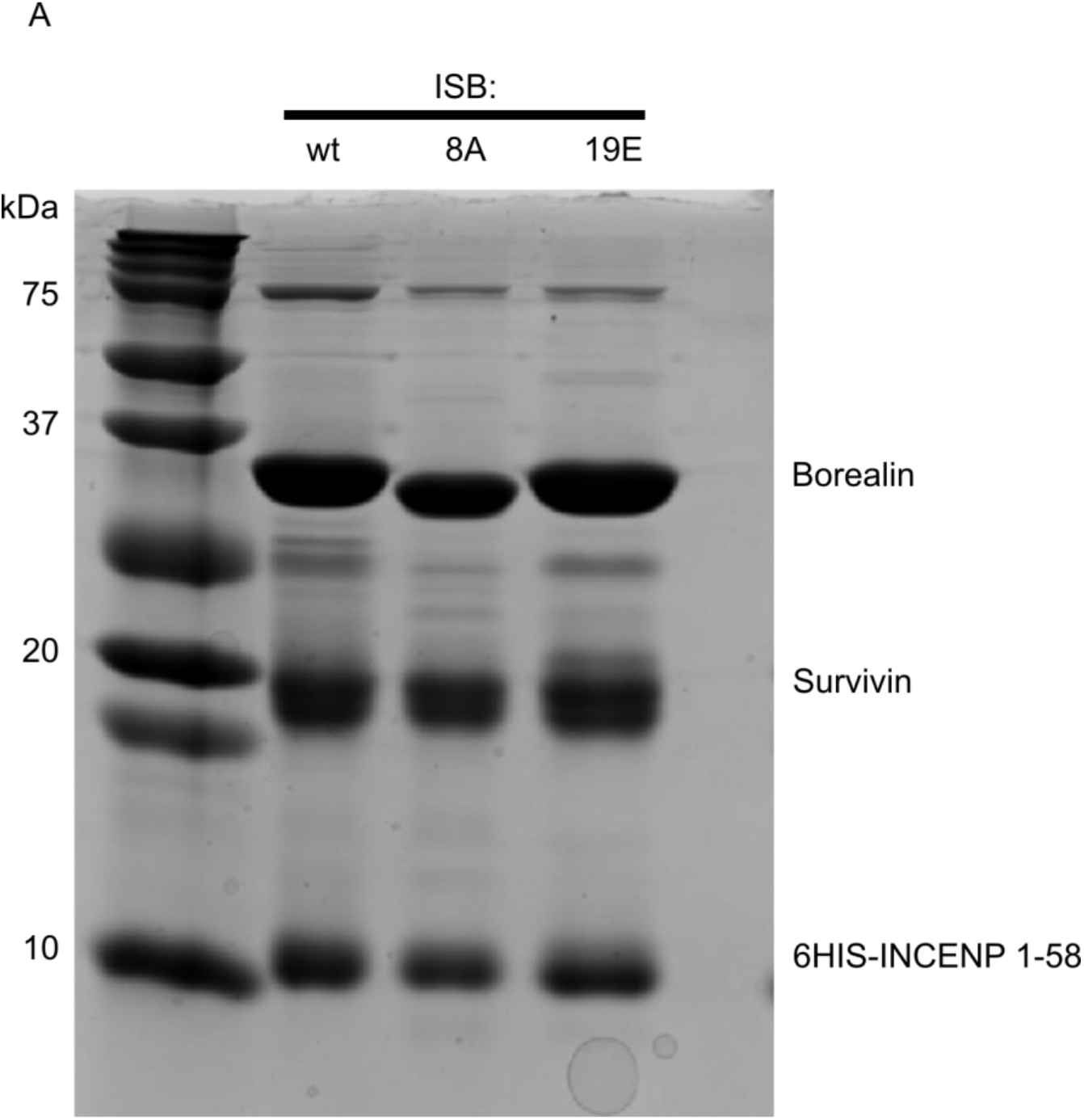
Purity of recombinant CEN-Borealin^wt^, CEN-Borealin^8A^, CEN-Borealin^R17E,R19E,R20E^ protein preps. A. SDS-PAGE gel stained with Coomassie to show components and purity of the CEN wild type and mutant preps used in MTs polymerization and bundling experiments.

**Supplementary Figure 6.**
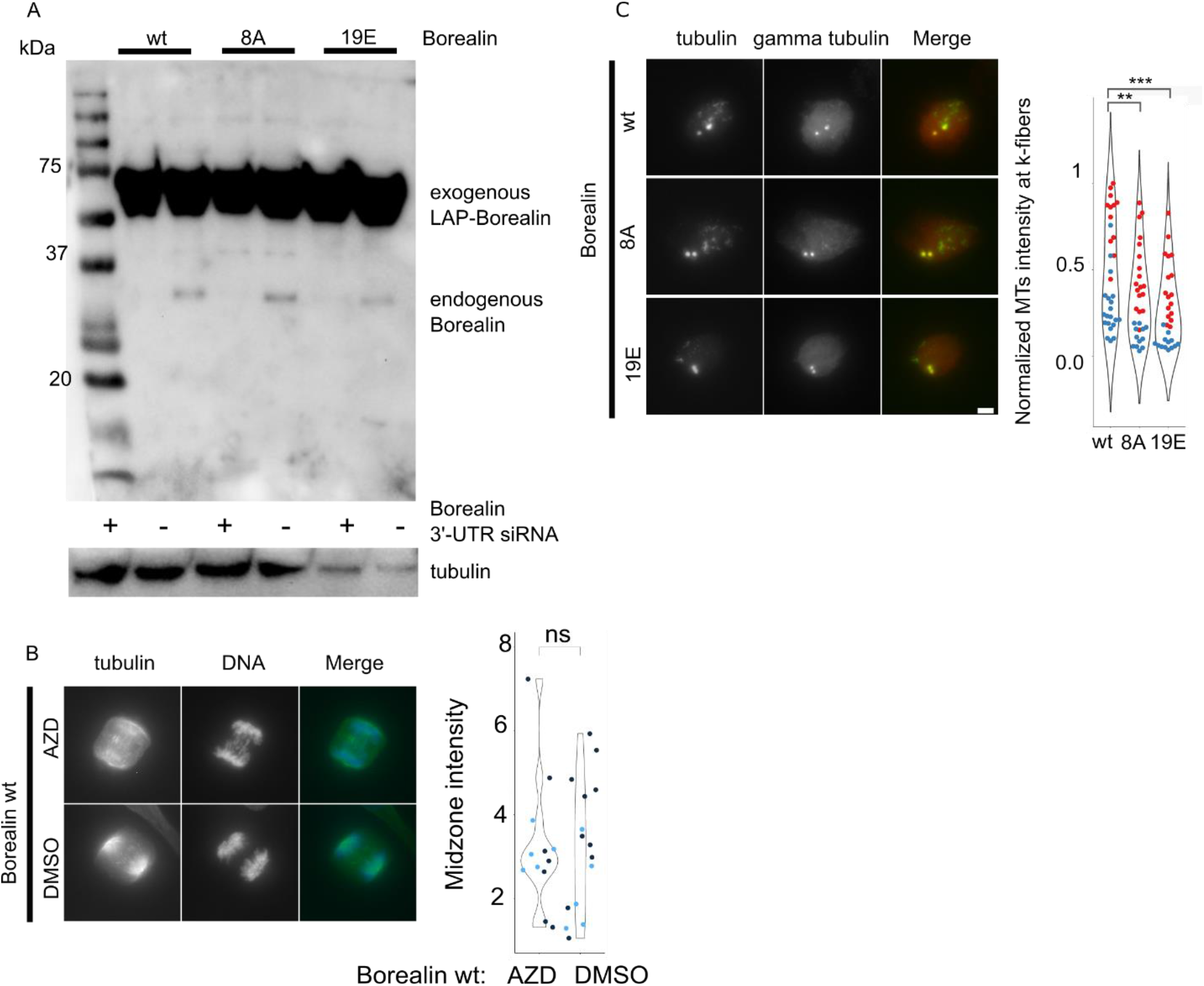
Western blot of Borealin siRNA knock-down and replacement and immunofluorescence analysis of Borealin depleted cells. A. Western blot showing Borealin and LAP-Borealin levels in HeLa cells in midzone characterization and nocodazole washout experiments. B. Midzone MTs levels in HeLa LAP-Borealin^wt^ cells treated with AZD1152-HQPA n=(10, 14) and DMSO n=(20, 10). Experiment was repeated twice. For statistical analysis Welch’s t-test with Bonferroni correction was applied; p-value^AZD-DMSO^ = (0.9, 0.174). * = p<0.05, *** = p < 0.01, *** = p<0.001, ns = p>0.05. C. Fluorescence images of HeLa cells with endogenous Borealin replaced with wild type or indicated Borealin mutants and stained for α-tubulin, γ-tubulin, ACA and DAPI. Experiment was repeated two times; CEN-Borealin^wt^ n=(12, 20); CEN-Borealin^8A^ n=(18, 12); CEN-Borealin^R17E,R19E,R20E^ n=(17, 13). p-value^wt-8A^ = (1.2·10^-6^, 0.0062); p-value^R17E,R19E,R20E^ = (7.2·10^-9^, 0.0009). For statistical analysis Welch’s t-test with Bonferroni correction was applied; * = p<0.05, ** = p < 0.01, *** = p<0.001

